# TOP-SECRETS enables Cas9 nucleases to discriminate SNVs outside of PAMs

**DOI:** 10.1101/2025.05.06.652491

**Authors:** Ashley Herring-Nicholas, Stephanie Fisher-Huynh, Eric A. Josephs

## Abstract

RNA-guided CRISPR nuclease Cas9 cannot reliably differentiate between single nucleotide variations (SNVs) of targeted DNA sequences determined by their guide RNA (gRNA): they typically exhibit similar nuclease activities at any of those variations, unless the variation occurs with specific sequence contexts known as protospacer adjacent motifs (PAMs). Our approach, “TOP-SECRETS,” generates gRNA variants that allow Cas9 ribonucleoproteins (RNPs) to reliably discriminate between healthy and disease-associated SNVs outside of PAMs.

## MAIN TEXT

Approximately 50% of human genetic variations known to be pathogenic consist of a single nucleotide variation (SNV) or point mutation.^2^ These include many important somatic mutations that are activating and autosomal-dominant: for example, simple missense mutations in the *KRAS* gene are among the most common driver mutations in many cancers, with the *KRAS*^G12D^ point mutation being the most common;^3^ and simple missense mutations in the *MED12* gene in codon 44 is linked to ∼50% of uterine leiomyomas, benign tumors that occur in about 60% -80% of women under 50 years of age and that are the most common cause of gynecologic surgery, with *MED12*^G44D^ being the most common.^4^ These examples of common, somatic, gain-of-function mutations might represent good candidates for targeted gene therapies where specific inactivation of only the disease allele but not the healthy one might result in a clinical benefit. In fact, the ability to consistently differentiate different SNVs using gene editing technologies would enable myriad new biotechnological applications, for example, specific targeting of SNPs for trait engineering of plant biotechnologies or for biomedical sciences. However, most gene editing biotechnologies, including CRISPR-Cas9 biotechnologies, lack the ability to discriminate between different SNVs or between healthy and pathogenic alleles that only differ by a single nucleotide^5^ outside of certain, specific genetic contexts known as a “protospacer adjacent motif” (PAM).^6^

That is, while CRISPR-Cas9 biotechnologies have revolutionized the ability to introduce targeted genetic mutation at specific DNA sequences determined by a modular 20-nt segment of the Cas9’s RNA co-factor known as their guide RNAs or gRNAs,^7^ under normal circumstances even they lack an ability to differentiate between SNVs. It is well-established that Cas9 ribonucleoproteins (RNPs) will exhibit nuclease activity and introduce double-strand breaks (DSBs) and mutations not only at their intended sequence target (their “protospacer”) (e.g., Figure 1Ai), but also at other “off-target” DNA sequences present in the host’s genome even if those sequences differ from the Cas9 RNP’s target by about 3 or 4 nts^8^ (e.g., Figure 1Aii) (and, biochemically, up to about 6 to 8 nts)^9^ of sequence divergence. While significant efforts have gone into limiting the effects of these “off-target” activities during therapeutic gene disruption by Cas9 – including the development of engineered, high-specificity variants of Cas9 such as “eCas9”^10^ – those engineered Cas9 variants are still not specific enough to be capable of discrimination between SNVs. The only exception is when the SNV occurs within a PAM:^5^ The CRISPR target-recognition process begins when the Cas9 enzyme itself first engages with a PAM sequence, which in the case of the biotechnologically-important Cas9 from *Streptococcus pyogenes* (SpyCas9) is ‘NGG’ (with N representing any nucleotide) and to a lesser extent ‘NAG’, positioned immediately 3’-of a protospacer sequence;^11^ the targeting segment of the gRNA known as the spacer is then able to base-pair with a complementary strand of the protospacer DNA, displacing the other strand, and when the spacer has stably base-paired entirely across its 20-nt length, the nuclease domains of the Cas9 enzyme shift into an active conformation to introduce strand breaks on the DNA molecules.^12^ In general, a Cas9 enzyme can only reliably differentiate SNVs if the variant results in the generation of a novel PAM sequence where there otherwise is not one, creating a new protospacer sequence that is not able to be recognized on the other allele; if the SNVs do not occur in PAM motifs, some other methods, such as introduction of additional mismatches into the spacer sequence, may also be attempted, but these typically result in significant attenuation of CRISPR activity at the desired target sequences as well. The requirement by Cas9 for a specific sequence context to differentiate SNVs represents a significant limitation of Cas9’s potential for the allele-specific targeting of pathogenic variants, simply because many pathogenic SNPs do not result in novel PAMs: in fact, in the examples listed above, both pathogenic mutations for *KRAS*^*G12D*^ (Figure 1Bii and 2A top) and *MED12*^G44D^ (Figure 2B bottom) occur in the healthy allele within ‘NGG’ sequence contexts, resulting in the conversion of a PAM sequence for *S. pyogenes* Cas9 in the healthy variant to a non-PAM sequence in the pathogenic variant — precisely the opposite conditions as required for current SNV-specific approaches using CRISPR based on the generation of a novel PAM.

**Figure 1.**
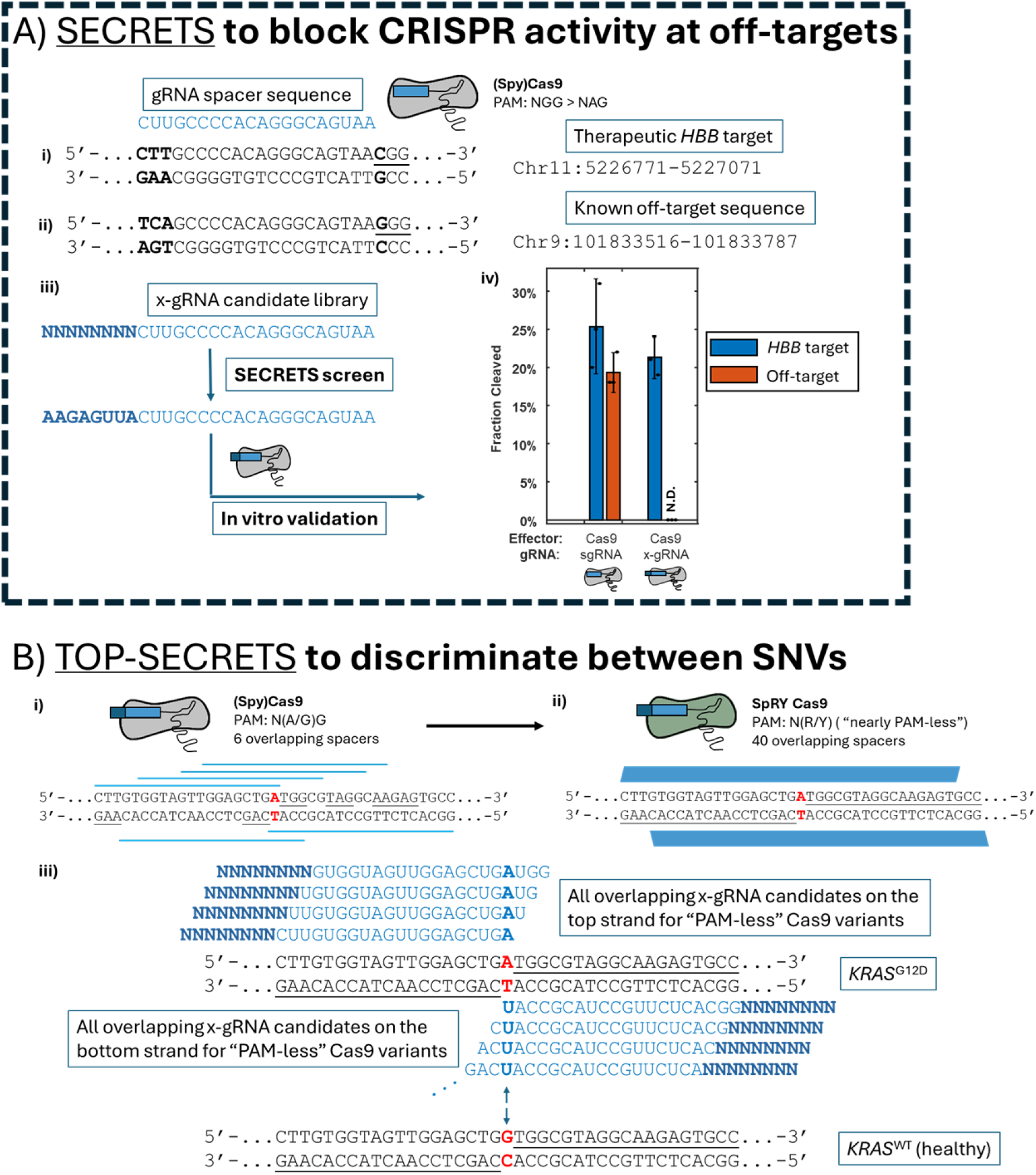
SECRETS and TOP-SECRETS effectively identify gRNAs with short 5’-extensions (x-gRNAs) capable of significant enhancements in Cas9 specificity (A) or their ability to discriminate between single nucleotide variants (SNV) of their targets (B), respectively. (A) During a SECRETS screen, gRNA variants for a target (spacer) of interest with every possible combination of short (8 nt) 5’-extensions (x-gRNA candidate library) in a process called SECRETS (Selection of Extended CRISPR RNAs with Enhanced Targeting and Specificity). x-gRNAs identified through SECRETS have been shown to eliminate nuclease activity by the Cas9 RNP at known off-targets while maintaining nuclease activity at their intended targets. Data in 1Aiv adapted from Ref. 1; dots represent independent experimental trials (n = 3), error bars are 95% confidence. N.D. = not detected. (B) In trying to discriminate a SNVs, (i) there are 6 possible 20 nt gRNA spacers (blue lines) for SpyCas9 – determined by the presence of PAM sequence recognized by SpyCas9, that is, a ‘NGG’ (N = any nucleotide) or a more-weakly recognized ‘NAG’ motif (underlined) – that overlap a pathogenic SNV (highlighted in red) and with which we can perform a SECRETS screen in combination (TOP-SECRETS). (ii) If TOP-SECRETS is performed with a Cas9 variant (SpRY Cas9) that recognizes spacers next to ‘NR’ (R = either purine) or ‘NY’ (Y = either pyrimidine), effectively rendering it “nearly PAM-less,” (iii) there are 20 possible spacers on the top strand and 20 spacers on the bottom strand (with potential “PAMs” underlined) that overlap the position of divergence between SNVs with which to perform TOP-SECRETS in combination (>2M possible x-gRNA variants).

**Figure 2.**
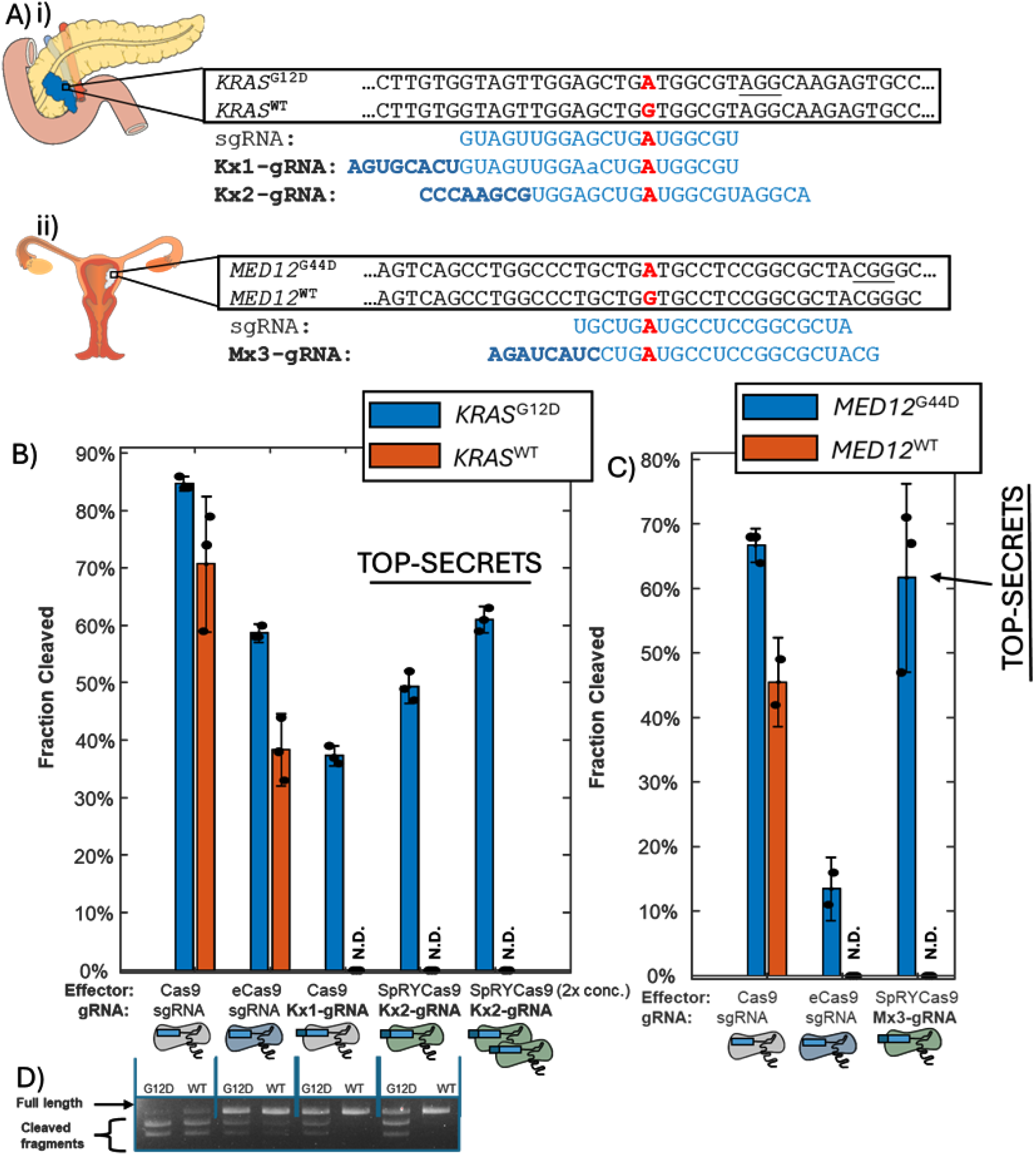
x-gRNAs identified through TOP-SECRETS can effectively discriminate between healthy and disease-associated SNVs of their targets. (A) The sequences and sequence contexts for a pathogenic Cas9 targets (KRAS^G12D^ mutation represented by pancreatic tumor (i) and MED12^G44D^ represented by a uterine fibroid (ii)) with their respective “healthy” variants (KRAS^WT^ and MED12^WT^) highlighted in red. A gRNA spacer and x-gRNA sequence determined by TOP-SECRETS are highlighted below, with the PAM for the target of the gRNA’s protospacer underlined. Lowercase represents mismatches with the target sequence. (B and C) Cas9 RNPs with x-gRNAs identified by TOP-SECRETS exhibit no nuclease activity at the healthy sequence variants of their pathogenic targets, even at 2x concentration relative to standard reaction conditions, and maintain activity at their pathogenic target better their even engineered “high-specificity” Cas9 variants (eCas9). Dots represent independent experimental trials (n = 2 or 3), error bars are 95% confidence. N.D. = not detected. (D) A representative gel electrophoresis assay comparing nuclease activity and specificity of Cas9 and eCas9 RNPs containing a standard gRNA for KRAS^G12D^ with Cas9 and SpRY Cas9(“near PAM-less”) RNPs containing x-gRNAs – despite SpRY Cas9 itself being known to exhibit attenuated activity compared to SpyCas9.

Previously, in attempting to address the problem of “off-target” activity by Cas9 RNPs, it was found that adding short (8 – 12 nucleotide) extensions to the 5’-end of the spacer sequence (Figures 1Aiii and 1Aiv), in such a way that these extra nucleotides were hypothesized to interact with the DNA-recognizing segment of the spacer, could result in orders-of-magnitude increases in targeting specificity at their intended target *vs*. at their known off-target sequences (that differed by 3 – 5 nucleotides or so);^13^ later, we developed a protocol, known as **S**election of **E**xtended **C**RISPR **R**NAs with **E**nhanced **T**argeting and **S**pecificity, or **SECRETS**, where we could screen hundreds of thousands of those gRNA variants known as extended gRNAs or x-gRNAs to identify extension sequences that resulted in Cas9 RNPs with improved activity and specificity profiles for a protospacer of interest, relative to its activity at known off-target sequences (Figure 1A).^1^ SECRETS works by screening all possible short nucleotide extensions (4^8^ or 65,536 possibilities in the example of 8 nucleotide extensions in Figure 1Aiii) to a spacer with both strong positive selection for x-gRNAs with high levels on-target activity and strong negative selection against activity at a specific known off-target sequences: using SECRETS, we have been able to consistently identify x-gRNAs capable of high levels of nuclease activity at their intended on-target sequence (activity consistent with their respective gRNA) while effectively abolishing activity at their “off-target” sequences (Figure 1Aiv).^1,14^

In pushing the limits of our SECRETS technique to determine if x-gRNAs could be used to discriminate between SNVs, we developed TOP-SECRETS (Figure 1B): where we hypothesized that if we **T**argeted all **O**verlapping **P**rotospacers (TOP) of a SNV while performing SECRETS screens (TOP-SECRETS), from this expanded search, we would be able to consistently identify x-gRNAs that can efficiently discriminate SNVs – even (and especially) if those SNVs were not located in PAM sequences. CRISPR enzymes are very sensitive not only to the 5’ extensions^1^ but also to where and of what species the improper nucleotide mismatch between the gRNA spacer and DNA target are;^15^ we hypothesized that by screening every possible overlapping spacer, we could identify x-gRNAs with exceptional specificity for those SNVs. Here, we show that TOP-SECRETS represents a robust methodology to screen millions of x-gRNA variants simultaneously and identify x-gRNAs that give a Cas9 RNP high levels of nuclease activity at multiple, disease-associated SNVs (including the two common ones mentioned above) but no detectable activity at their respective heathy variants when the SNV is located outside of PAM sequences. We expect that a “drop-in” technology — by a simple substitution of a gRNA with an x-gRNA — to enable Cas9 RNPs to readily differentiate SNVs in any sequence context, even outside of PAMs, will provide new opportunities for gene therapies by enable new CRISPR-based treatments for inherited and somatic genetic disorders and expand spectrum of applications of CRISPR biotechnologies that require this level of precision.

We first tested SECRETS to determine whether we could identify x-gRNAs capable of discriminating between pathogenic *KRAS*^*G12D*^ and healthy *KRAS*^*WT*^ sequences (Figure 1B and 2Ai). Using a standard single gRNA (sgRNA), both Cas9 and eCas9 were incapable of discriminating between the SNVs: Cas9 with a sgRNA targeting the *KRAS*^*G12D*^ sequence still exhibited 83% of its on-target cleavage activity at a *KRAS*^*WT*^ sequence, with eCas9 faring only moderately better with 65% relative activity (Figures 2B and S1). Using SECRETS, we performed screens for x-gRNA libraries of spacers with 8 nucleotide extensions for that protospacer; from the approximately 65,000 possible x-gRNAs for that protospacer overall, we were able to identify an x-gRNA that still allowed the Cas9 RNP to maintain activity at the *KRAS*^*G12D*^ (∼40%) but with no detectable activity “off-target” at the wild-type *KRAS*^*WT*^ sequence (Figures 2B and S1), demonstrating that we could identify x-gRNAs with strong SNV-discriminating capabilities.

However, SpyCas9 has only 6 possible protospacers that overlap the site of the *KRAS*^*G12D*^ SNV, determined by the presence of ‘NGG’ or (weaker but still recognized) ‘NAG’ PAM sites within 20 nt (Figure 1Bi). To see if we could further SNV-discriminating x-gRNAs with improved on-target activities, we decided to perform TOP-SECRETS by screening all possible overlapping protospacer sequences for a Cas9 enzyme that had been engineered to have relaxed PAM requirements (Figure 1Bii):^16^ SpRY Cas9, which has a slightly greater level of activity with PAM ‘NR’ (where R = A or G) over PAM ‘NY’ (where Y = C or T) but with high levels of activity at targets with both PAMs, rendering it effectively “near PAM-less”. Compared to the SpyCas9, which had 6 potential protospacers that overlapped the *KRAS*^*G12D*^ mutations (next to NGG or alternative PAM NAG), relaxing the PAM requirements for the Cas9 nuclease allowed us to test all possible spacer sequences for any potential protospacer that overlapped the *KRAS*^*G12D*^ SNV on the top and bottom strands (20 on the top and 20 on the bottom): in principle, >2.6M x-gRNA candidates (Figure 1Biii). After TOP-SECRETS, we were able to identify an x-gRNA that in vitro exhibited >70% activity of the Cas9 and standard sgRNA at the *KRAS*^*G12D*^ sequence — in other words, exhibiting the same “on-target” nuclease activity as eCas9 — and that exhibited no detectable activity at the *KRAS*^*WT*^ sequence (Figures 2B and 2D) – even when using twice the concentration of Cas9 RNP. We subsequently repeated TOP-SECRETS using SpRY Cas9 and targeting the pathogenic *MED12*^G44D^ SNV: while eCas9 suffered significant attenuation of activity on-target (∼20% activity), SpRY Cas9 with the x-gRNA identified by TOP-SECRETS exhibited as high a level of activity at the *MED12*^G44D^ SNV as the standard SpyCas9 (p = 0.5958, one-sided t-test, n = 3) and yet no detectable nuclease activity at the healthy *MED12*^WT^ sequence (Figure 2C). These findings demonstrate that TOP-SECRETS can effectively identify x-gRNAs with the same level of activity as normal CRISPR approaches while still exhibiting extreme SNV discrimination even when the SNV of interest does not generate a novel PAM sequence.

We also note that these comparable levels of on-target activity at pathogenic SNVs (with no detectable cleavage activity at healthy SNVs) with regular CRISPR approaches using Cas9 and eCas9 are particularly remarkable considering that SpRY Cas9 has been found to be relatively inefficient for gene editing applications compared to SpyCas9,^17^ as its lack of PAM requirement effectively reduces its rate of target recognition and cleavage. Since PAM recognition is believed to be structurally uncoupled from the DNA target recognition and nuclease domains of the Cas9 enzyme,^17,18^ we hypothesized that x-gRNAs identified using TOP-SECRETS with SpRY Cas9 (to screen every possible overlapping protospacer for SNV) would be even more effective if introduced back into Cas9 enzymes that required PAM binding to initiate target recognition; we predicted this would have the effect of both increasing on-target activity, and reducing the potential for any (genome-wide) off-target interactions. The most effective x-gRNA we identified for *KRAS*^*G12D*^ from TOP-SECRETS, which we called Kx2-gRNA, has a protospacer sequence immediately upstream of a ‘AGAG’ sequence. Engineered SpyCas9 variants Cas9^*EQR*^ and Cas9^*VQR*^ both recognize ‘NGAG’ motifs as PAMs (Figure 3A);^11,18,19^ so we then compared those Cas9 variants with SpRY Cas9 RNPs using Kx2-gRNA during an *in vitro* assay of both nuclease activity on-target and specificity (Figures 3B and S2).^1^ We found that, as expected, the extension sequence of x-gRNA necessary for specificity – those using the respective sgRNA of Kx2-gRNA performed orders of magnitude worse in the activity / specificity survival assay than those using the Kx2-gRNA – but we also found that Cas9^*EQR*^ and Cas9^*VQR*^ RNPs both performed better than SpRY Cas9 RNPs. These findings suggest that x-gRNAs discovered using “near PAM-less” SpRY Cas9 are not only still functional in other Cas9 variants but that they can be even more effective when re-introduced back into Cas9 variants with more stringent PAMs for the purpose of CRISPR-based discrimination of SNVs outside of a PAM sequence.

**Figure 3.**
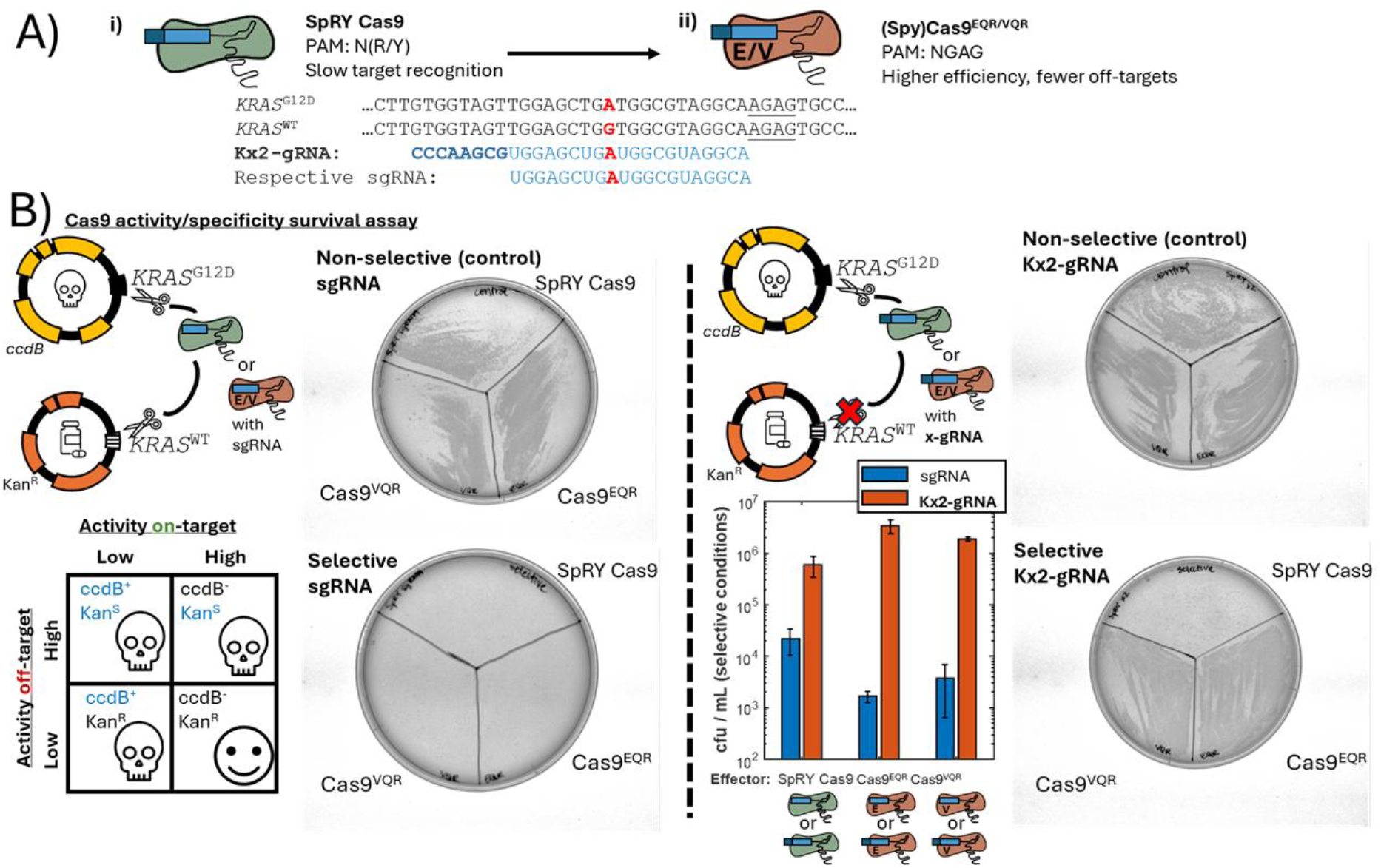
SNV-discriminating x-gRNAs identified from TOP-SECRETS with “PAM-less” Cas9 variants are even more effective in Cas9 variants with more stringent PAMs. A) While SpRY Cas9 was used to discover SNV-discriminating x-gRNAs for KRAS^G12D^ with TOP-SECRETS and the identified x-gRNA (Kx2-gRNA) targets a sequence with a PAM (NGAG, underlined) that is not recognized by the standard SpyCas9, that PAM is instead recognized by Cas9 variants Cas9^EQR^ and Cas9^VQR^. Below, the respective sgRNA for Kx2-gRNA that does not have the 5’-extension sequence. B) By performing a Cas9 activity/specificity survival assay, where bacterial cells can only survive if they have a Cas9 (or Cas9 variant) RNP that exhibits strong activity at the pathogenic KRAS target sequence and minimized / no activity at the healthy variation of the KRAS sequence, we find: not only is the extension sequence of the x-gRNA necessary for in vitro SNV-discrimination, but that Cas9 variants with more stringent PAMs (Cas9^EQR^ and Cas9^VQR^) are even more effective in their SNV-discriminating specificity and activity than “near PAM-less” SpRY Cas9. cfu = colony forming units. n = 3 independent trials, error bars are 95% confidence.

On multiple occasions, x-gRNAs have been shown to be stable, functional, and capable of increasing specificity with Cas9 in human cell lines for targeted mutagenesis;^1,13^ while each SNV-specific x-gRNA generated by TOP-SECRETS will need to be tested for mutational efficiency in the cell of interest (like any other gRNA), here we have shown a robust methodology to efficiently identify x-gRNAs that give Cas9 the ability to differentiate SNVs outside of PAM sequences in a “design-free” manner and without “trial and error.” The addition of extra 5’-nucleotides in the x-gRNAs were also previously found not to increase the number of potential “off-target” sites genome-wide; this finding is likely a result of their being outside of the DNA-targeting segment / spacer of the gRNA.^1,13^ We note again that x-gRNAs identified using TOP-SECRETS represent a “drop-in” technology — they can simply be substituted for the standard gRNA for whatever application of Cas9 is being performed, with the only difference being the addition of a few (here, 8) extra nucleotides to the 5’-end.

Ultimately, we expect that the ability to generate SNV-specific Cas9 RNPs (Figure 4) will enable a number of new biotechnological applications, including allele-specific gene editing as well as other CRISPR-based modalities^7^ such as base editing or prime editing, and improve on other non-therapeutic applications of CRISPR such as biosensing, trait engineering for plant biotechnologies, or biomedical research. SNV-specific gene editing can also in principle increase the safety of gene therapies for somatic genetic disorders by limiting the range of nuclease activity only to tissue with disease-causing variants. Furthermore, an efficient ability to recognize SNVs with Cas9 nucleases using a technique like TOP-SECRETS may help to make CRISPR-based treatments more broadly accessible to those with genetic disorders caused by uncommon SNVs that might occur outside of PAM sequences.^20^

**Figure 4.**
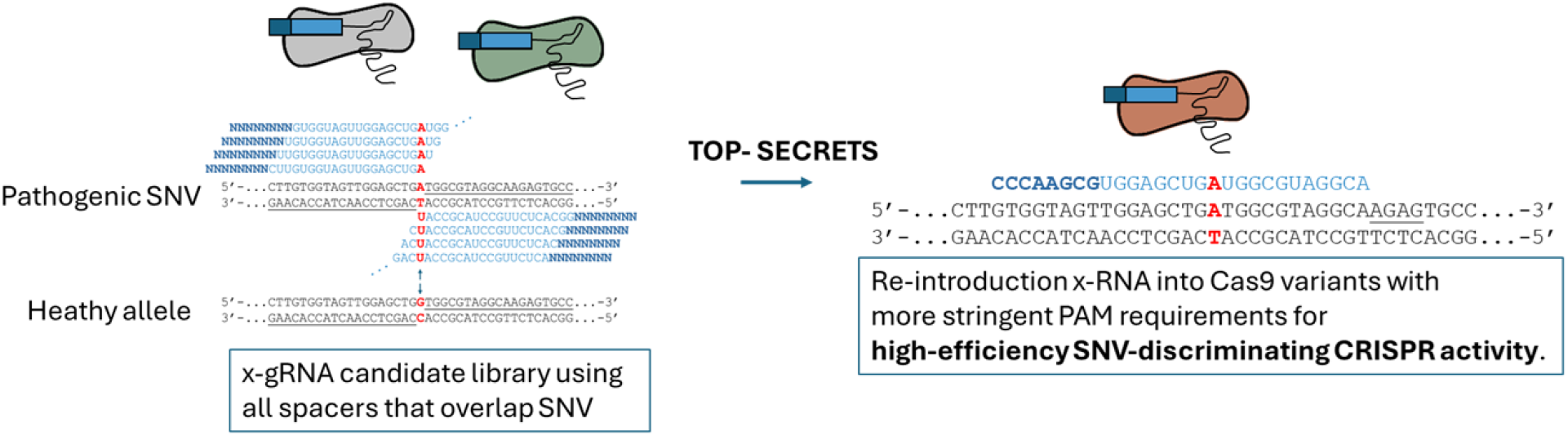
Summary of TOP-SECRETS to generate SNV-specific CRISPR RNPs for SNVs outside of PAMs. After a SNV of interest is identified, a randomized library of x-gRNA candidates with every spacer sequence that Targets every possible Overlapping Protospacer (TOP) containing the SNV are generated and screened using the SECRETS protocol. When SNV-specific x-gRNA candidates are identified and validated, if the sequence next to the protospacer for that x-gRNA is not the canonical ‘NGG’ motif for SpyCas9, those x-gRNAs can still be re re-introduced into SpyCas9 variants (Cas9^EQR^, for example) the sequence adjacent to the protospacer that recognize as a PAM to increase nuclease efficiency and activity while maintaining SNV discrimination.

## Materials and Methods

### DNA oligonucleotides, dsDNA, and plasmids

DNA sequences for all oligonucleotides and DNA fragments are listed in Supplementary Table 1.

### E. coli strains

All cloning was performed using New England Biolabs (NEB) 10-beta cells (NEB #C3020K) or 5-alpha cells (NEB #C2987H), and all TOP-SECRETS assays were performed in Stbl2 cells (Invitrogen #10268019), grown at 30°C.

### Cloning SECRETS plasmids and x-gRNA libraries: Plasmids

The cloning of plasmids for SECRETS and TOP-SECRETS was performed similarly previously described in Ref. 1 with some minor changes. Briefly: plasmid pSECRETS-A (Addgene #196986) contains an chloramphenicol resistance cassette and a gene for tetracycline-inducible expression of Cas9. To generate a modified pSECRETS-A with SpRY Cas9, the gene for SpRY Cas9 was PCR amplified from the pET-SpRY-NLS-6xHis (gbr2101) plasmid (Addgene #181743) and combined with a PCR’ed segment of pSECRETS-A (amplified with a different primer set; Supplementary Table 1) *via* HiFi Assembly. The pET28-SpRY-NLS-6xHis (gbr2101) was a gift from Benjamin Kleinstiver (Addgene plasmid #181743; http://n2t.net/addgene:181743; RRID: Addgene_181743).

To clone the x-gRNA libraries into pSECRETS-B (Addgene Plasmid #196987), a low copy number plasmid containing a kanamycin resistance cassette, for SECRETS, x-gRNA libraries containing a single spacer sequence and randomized 8 nt sequences (‘NNNNNNNN’) along with the “off-target” sequence (the “healthy” SNV, along with the 34 nucleotides flanking on each side) were cloned and maintained as previously described. For TOP-SECRETS, x-gRNA libraries containing all spacer sequences that overlapped the SNV of interest transformations were done as a series of 9 replicates to ensure coverage of the larger library set (now >2.6M x-gRNAs in the case of SpRY Cas9-based TOP-SECRETS).

To generate into pSECRETS-C, which is a derivative of p11-LacY-wtx1 (Addgene Plasmid #69056), a high-copy number plasmid containing an arabinose-inducible (glucose-repressed) expression of toxin ccdB, an ampiciliin resistance cassette, and the on-target sequence (the “pathogenic” SNV, along with the 34 nucleotides flanking on each side), the on-target sequence was cloned into p11-LacY-wtx1 as previously described.

### The SECRETS and TOP-SECRETS protocol and analysis

*Selection of extended g-RNAs (SECRETS protocol) and Targeting all Overlapping Protospacers-SECRETS (TOP-SECRETS)*

Once the correct plasmids were cloned, TOP-SECRETS was performed with a similar protocol as the SECRETS assay described in Ref. 1. Briefly, Stbl2 *E. coli* cells were electroporated with Cas9 or SpRY-Cas9 pSECRETS-A plasmids (50 ng DNA) and recovered in 950 uL of 10-beta/Stable outgrowth media (NEB #B9035S) at 37°C for 1 hour before being plated on LB+chloramphenicol plates. These pSECRETS-A expressing cells were made electrocompetent then electroporated with the desired pSECRETS-C plasmid (50 ng DNA) and recovered in LB+gluc (0.1% f.c.), before being plated on selective plates at 30°C for 24 hours. Colonies were sequenced for pSECRETS-C propagation and once confirmed, these cells were made electrocompetent for pSECRETS-B uptake and screening. Stbl2 *E. coli* cells now expressing pSECRETS-A and -C were electroporated with their corresponding pSECRETS-B plasmid (100 ng DNA) containing the off-target site and sgRNA (or x-gRNA libraries). Following recovery (30°C for 75 mins), cells were centrifuged, and supernatant was replaced with fresh LB before inoculating 0.5 mL of the culture into 7 mL liquid LB for selective or non-selective conditions and grown overnight. After miniprep (NEB #T1010L) of the resulting cultures, samples were PCR’ed across the gRNA segment and prepared for Illumina next-generation sequencing using variants of the following primers:

SECRETS-BSeq5 `5’-[NGS adapter][barcode]-GAGCGGATACATATTTGAATG-3’`

SECRETS-BSeq3 `5’-[NGS adapter][barcode]-AAGTTGATAACGGACTAGCC-3’`

Biological replicates were performed in triplicate and technical replicates were performed in duplicate.

### Analysis of SECRETS outcomes

Small-scale next-generation sequencing of samples from the SECRETS assay was performed (Amplicon-EZ, Azenta Inc.; at least 50,000 reads, typically >300,000). Custom code was written in MATLAB (Mathworks; Natick, MA) to extract and count the 5’-extensions from the x-gRNA sequence of each read; however, in principle, a short line of code can be written to the same effect following the approach found in Ref. 21. The number of unique 5’-extensions was enumerated per sample, normalized to the total number of reads per sample, then averaged across technical replicates (n = 2). The normalized number of reads per 5’-extension was then sorted from most prevalent to least, and the top five most prevalent 5’-extensions per gRNA were selected for further characterization.

### *In vitro* validation of x-gRNAs

#### Cas9 ribonucleoprotein (RNP) generation

DNA oligos of sgRNAs and x-gRNAs were designed according to the EnGen sgRNA Synthesis Kit (NEB #E3322) to add 5’-T7 RNA polymerase promoter sequence and 3’-Cas9 crRNA sequence and purchased from IDT and resuspended to a stock concentration of 100 μM. If the (x-)gRNA did not have an initial 5’-G necessary for efficient T7 RNA polymerase transcription, one was added to the DNA oligo sequence. For sgRNA synthesis, oligos were diluted 100x (1 μM) for use with the EnGen sgRNA Synthesis Kit, per manufacturer’s instructions. Cas9 RNPs were formed following the IDT Alt-R CRISPR-Cas9 System – *In vitro* cleavage of target DNA with ribonucleoprotein complex protocol (Option 2). Cas9 enzyme (Sigma Aldrich, #CAS9PROT-250UG), eCas9 enzyme (Sigma Aldrich #ESPCAS9PRO-50UG), or SpRY Cas9 enzyme (NEB, #M0669T) and (x-)gRNA were combined in equimolar amounts in dilution buffer (supplied with the (e)Cas9 proteins from Sigma) and incubated at room temperature for 10 minutes. Following incubation, RNPs were stored at -80°C or immediately used for *in vitro* digestion reactions.

#### *In vitro* digestion reactions

DNA targets (300 bp long) containing the target sequence (located ∼200 bp from the end) and the flanking genomic context were synthesized by Twist Bioscience, PCR amplified using the provided universal primers, purified, and resuspended in nuclease-free water to 100 nM. Three technical replications of reactions were assembled in the following order: 7 μL nuclease-free water, 1 μL 10x Cas9 Nuclease Reaction buffer (200 mM HEPES, 1 M NaCl, 50 mM MgCl2, 1 mM EDTA (pH 6.5 at 25°C)), 1 μL target DNA substrate (100 nM), and 1 μL Cas9-RNP (1 μM), then incubated for 1 hour at 37°C followed by 1 μL proteinase K and 1 μL RNAse A digestion at 56°C for 10 minutes (NEB #P8107S and #T3018L, respectively). Products were resolved on a 2% agarose gel stained with SYBR Gold, and the fraction cleaved was analyzed using ImageJ. The band fluorescence was measured, normalized based on the DNA fragment length, and the percentage cleaved was determined using the following equation:

[sum of cleaved band intensities/(sum of cleaved and parental band intensities)]x100%

### *Cas9 RNP activity / specificity survival assay (in* E. coli)

#### Plasmids

To generate pSECRETS-A derivatives that expressed Cas9^EQR^ and Cas9^VQR^, pSECRETS-A was used as the template along with the appropriate primers to introduce the Cas9^EQR^ or Cas9^VQR^ mutations (D1135E/R1335Q/T1337R or D1135V/R1335Q/T1337R; Supplementary Table 1) into the SpyCas9 gene via HiFi assembly. A pSECRETS-B derivative containing the *KRAS*^WT^ sequence (the “off-target”) and expression cassette for Kx2-gRNA or its respective sgRNA (same spacer sequence but no extension) was generated as described above. The pSECRETS-C plasmid contained the *KRAS*^G12D^ target sequence along with its 15-bp genomic context was generated as described above.

#### Survival assay

After cloning, colonies were sequenced for the correct sequence incorporation before simultaneously electroporating SpRY (or Cas9^EQR^ and Cas9^VQR^)-containing derivatives of pSECRETS-A and pSECRETS-C plasmid containing the *KRAS*^G12D^ target sequence (50 ng each) into Stbl2 *E. coli* cells. Cells were recovered in 10-beta/Stable outgrowth media (NEB # B9035S) for 1 hour at 37°C, then plated on LB+chlor+amp plates. Colonies were made competent, then electroporated with *KRAS*^G12D^ sgRNA (or Kx2-gRNA) containing pSECRETS-B (50 ng DNA). For the assay, cells were recovered in 1 mL of LB w/ glucose and aTc (0.1%, 10 ng/mL; f.c., respectively) at 30°C for 75 minutes, then pelleted (6,000 x g, 5 mins). The supernatant was discarded, and the cells were resuspended in 0.5 mL fresh LB before plating them under selective conditions. For cfu/mL calculations, the recovery was diluted and 10 uL drops were dispensed on selective plates (as in Figure S2).

#### Statistics and reproducibility

All samples were the result of at least 3 biological and 2 technical and biological replicates. Statistical analysis such as two-sided T-tests and generation of confidence intervals were performed using MATLAB or Microsoft Excel.

## Data availability

Source data and next-generation sequencing data obtained in these experiments will be made publicly available at the appropriate repositories after peer review and publication of the manuscript or upon request. All other data are available from the corresponding author upon reasonable request.

## Code availability

Code to perform and analyze the results of TOP-SECRETS will be made available after peer review and publication of the manuscript or upon reasonable request.

## Materials availability

Materials to perform TOP-SECRETS will be made available after peer review and publication of the manuscript or upon reasonable request.

## Acknowledgements

We thank members of the Josephs laboratory for careful reading of the manuscript. The work was funded by grants from the National Institute of General Medical Sciences [R35GM133483 to EAJ and T34GM149494 to SFH] and the National Institute of Biomedical Imaging and Bioengineering [R21EB033595 to EAJ] of the US National Institutes of Health. The content is solely the responsibility of the authors and does not necessarily represent the official views of the National Institutes of Health. The work was also supported by funds from the SUNY Empire Innovation Program and departmental funds from the Department of Biomedical Engineering at Stony Brook University. AHN received support as a UNC Greensboro Minerva Scholar and ICONS Scholar [US Department of Defense Contract #W911QY2220006]. This work in part was performed in part at the Joint School of Nanoscience and Nanoengineering of UNC Greensboro and North Carolina Agricultural and Technical State University, a member of the Southeastern Nanotechnology Infrastructure Corridor (SENIC) and National Nanotechnology Coordinated Infrastructure (NNCI), which is supported by the National Science Foundation [Grant ECCS-1542174].

## Declaration of interests

AHN and EAJ are inventors on a provisional patent filed by the UNC Greensboro for related technologies. AHN and EAJ are inventors on another patent application related to SECRETS and EAJ is an inventor on other patents and patent applications related to CRISPR technologies.

