## Supplementary Information for "TOP-SECRETS enables Cas9 nucleases to discriminate SNVs outside of PAMs"

### SUPPLEMENTARY MATERIAL

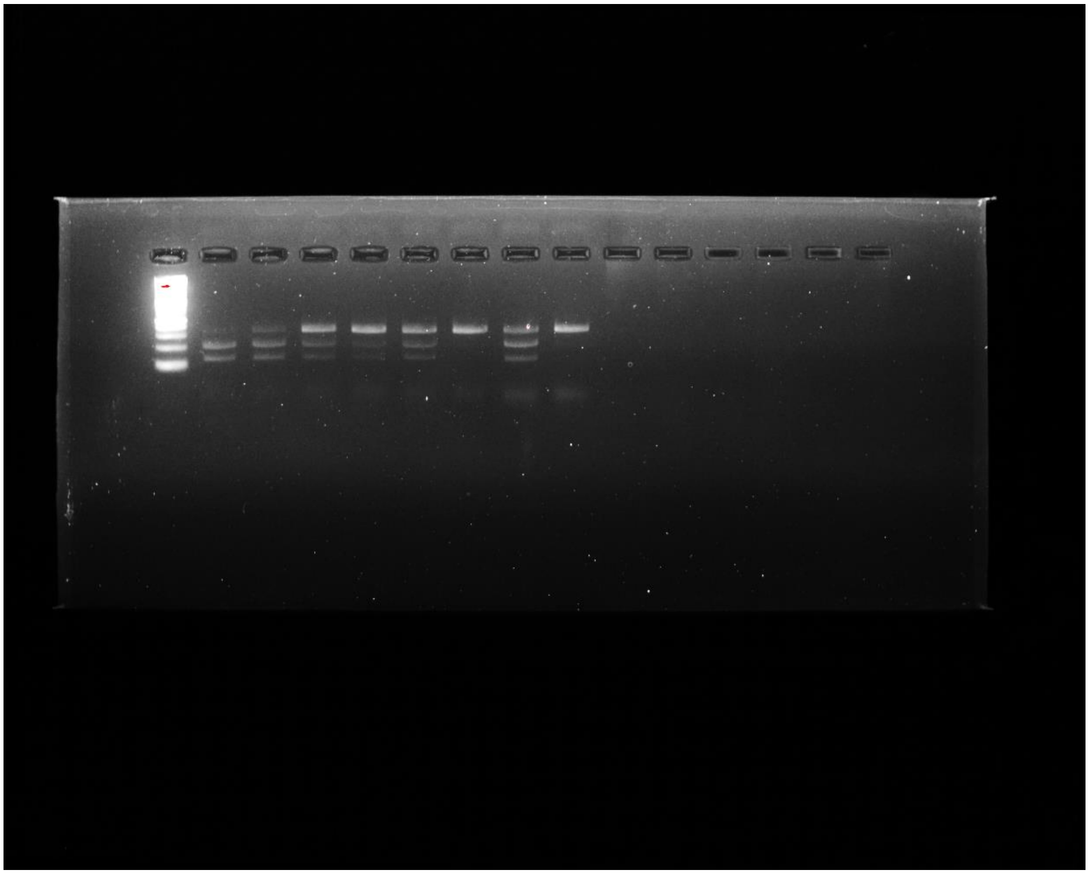

**Figure S1.** Uncropped gel image from Figure 2D. Lanes, left to right are: (1) DNA ladder; (2) SpyCas9 with sgRNA with 300 bp *KRAS*$^{G12D}$ target; (3) SpyCas9 with sgRNA with 300 bp *KRAS*$^{WT}$ target; (4) eCas9 with sgRNA with 300 bp *KRAS*$^{G12D}$ target; (5) eCas9 with sgRNA with 300 bp *KRAS*$^{WT}$ target; (6) SpyCas9 with Kx1-gRNA with 300 bp *KRAS*$^{G12D}$ target; (7) SpyCas9 with Kx1-gRNA with 300 bp *KRAS*$^{WT}$ target; (8) SpRY Cas9 with Kx2-gRNA with 300 bp *KRAS*$^{G12D}$ target; (9) SpRY Cas9 with sgRNA with 300 bp *KRAS*$^{WT}$ target.

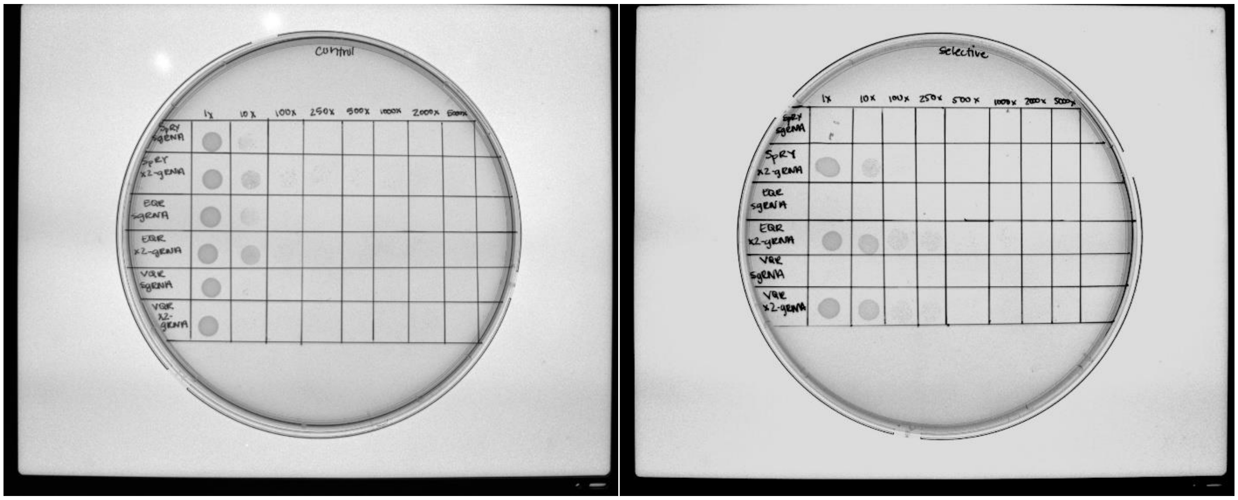


**Figure S2.** Example results of the in vitro specificity / activity bacterial survival assay, from Figure 3B. *E. coli* containing (1) a plasmid containing a cassette for inducible expression of SpRY Cas9, SpyCas9$^{EQR}$, or SpyCas9$^{VQR}$; (2) a high-copy number plasmid containing both the *KRAS*$^{G12}$ sequence (+ flanking 15 bp) and cassette for inducible expression of toxin ccdB; and (3) a low-copy number plasmid containing both the *KRAS*$^{WT}$ sequence (+ flanking 15 bp) and cassette for inducible expression of the KRAS sgRNA or Kx1-gRNA (shown in Figure 3A) and kanamycin resistance cassette.

(left) Under control conditions, both the ccdB expression cassette is held in respressive conditions and the plates do not contain the antibiotic kanamycin. After transfection of the gRNA and ccdB plasmid, 10 uL of bacteria are plated at differing levels of dilution until single colonies are able to be unambiguously counted. (right) Under selective conditions, both the ccdB expression cassette is in inducing contions for expression and the plates contain the antibiotic kanamycin: only bacteria where the Cas9 variant RNP that exhibit high levels of activity to introduce DSBs into high-copy toxin plasmid with the *KRAS*$^{G12}$ plasmid and very low levels of activity to prevent introducing DSBs into the low copy kanamycin resistance plasmid are able to survive the screen.

**Supplementary Table 1. Oligonucleotide and primer sequences**

| Name | Sequence | Note |
|---|---|---|
| pBbA2K-RFP-Fwd | AGTCACACTGGCTCACCTTC | Cloning (pSECRETS-A) |
| pBbA2K-RFP-Rev | TAACAACCCGTAAACTCGCC | Cloning (pSECRETS-A) |
| Cas9-Fwd | GCACAAATAGCGTCGGATGG | Cloning (pSECRETS-A) |
| Cas9-Rev | GAAGGTGAGCCAGTGTGACT | Cloning (pSECRETS-A) |
| SpRY-Fwd | CTCTTTTATTTGACAGTGGAGAGACCGCTGAGCGTACTC | Cloning (pSECRETS-A) |
| SpRY-Rev | CCTGGAGATCCTTACTCGAGTTAGTCCCCGCCTAACTG | Cloning (pSECRETS-A) |
| pBbA2K-SpRY-Fwd | CTCGAGTAAGGATCTCCAG | Cloning (pSECRETS-A) |
| pBbA2K-SpRY-Rev | TCCACTGTCAAATAAAAGAG | Cloning (pSECRETS-A) |
| SECRETS-Fwd-gRNA | CACTGCTTACTGGCTTATCG | Cloning (pSECRETS-B) |
| SECRETS-Rev-gRNA | CTTGCTATTTCTAGCTCTAAAAC | Cloning (pSECRETS-B) |
| x-gRNA library | CCACTGCTTACTGGCTTATCGGAAG [OFF-TARGET SEQUENCE, 20 NUCLEOTIDE + 3 NUCLEOTIDE PAM] TTTCCCTATCAGTGATAGAGATTGACATCCCTATCAGTGATAGAG ATACTGAGCAC [5'- EXTENSION LIBRARY (WHEN APPLICABLE): NNNNNNNN, 20 NUCLEOTIDE SPACER SEQUENCE] GTTTTAGAGCTAGAAATAGCAAG TTTCCCTATCAGTGATAGAGATTGACATCCCTATCAGTGATAGAG ATACTGAGCAC | Cloning (pSECRETS-B) |
| pSECRETS-B-Fwd | CGATAAGCCAGTAAGCAGTG | Cloning (pSECRETS-B) |
| pSECRETS-B-Rev | GTTTTAGAGCTAGAAATAGCAAG | Cloning (pSECRETS-B) |
| pSECRETS-C-Fwd | AAGCTTGGCTGTTTTGGCG | Cloning (pSECRETS-C) |
| pSECRETS-C-Rev | GCGTGATATTACCCTGTTATCCC | Cloning (pSECRETS-C) |
| pSECRETS -C-targets | ATAACAGGGTAATATCACGC-[15 BP UPSTREAM GENOMIC SEQUENCE CONTEXT] + [20 BP TARGET SEQUENCE + 3 BP PAM] + [15 BP DOWNSTREAM GENOMIC SEQUENCE CONTEXT] AAGCTTGGCTGTTTTGGCGG | Cloning (pSECRETS-C) |
| SECRETS-BSeq51 | ACACTCTTTCCCTACACGACGCTCTTCCGATCT TTTT GAGCGGATACATATTTGAATG | Sequencing pSECRETS-B (x-)gRNA libraries |
| SECRETS-BSeq52 | ACACTCTTTCCCTACACGACGCTCTTCCGATCT AAAA A GAGCGGATACATATTTGAATG | Sequencing pSECRETS-B (x-)gRNA libraries |
| SECRETS-BSeq53 | ACACTCTTTCCCTACACGACGCTCTTCCGATCT GGGG TA GAGCGGATACATATTTGAATG | Sequencing pSECRETS-B (x-)gRNA libraries |
| SECRETS-BSeq54 | ACACTCTTTCCCTACACGACGCTCTTCCGATCT CCCC GCC GAGCGGATACATATTTGAATG | Sequencing pSECRETS-B (x-)gRNA libraries |

| | | |
|---|---|---|
| SECRETS-BSeq31 | GACTGGAGTTCAGACGTGTGCTCTTCCGATCT TTTT AAGTTGATAACGGACTAGCC | Sequencing pSECRETS-B (x-)gRNA libraries |
| SECRETS-BSeq32 | GACTGGAGTTCAGACGTGTGCTCTTCCGATCT AAAA A AAGTTGATAACGGACTAGCC | Sequencing pSECRETS-B (x-)gRNA libraries |
| SECRETS-BSeq33 | GACTGGAGTTCAGACGTGTGCTCTTCCGATCT GGGG TA AAGTTGATAACGGACTAGCC | Sequencing pSECRETS-B (x-)gRNA libraries |
| SECRETS-BSeq34 | GACTGGAGTTCAGACGTGTGCTCTTCCGATCT CCCC GCC AAGTTGATAACGGACTAGCC | Sequencing pSECRETS-B (x-)gRNA libraries |
| QR-1-Fwd | TATTTTGATACAACAATTGATCGTAAACAATATAGGTCTACAAAAGAAGTTTTAGATGCC | Cloning (E/VQR vector) |
| E-1-Rev | CCACTAGGACTGAATAAGCTACCGTTGGACTTTCAAAACCACCATATTTTTTTGGATCCC | Cloning (EQR vector) |
| V-1-Rev | CCACTAGGACTGAATAAGCTACCGTTGGACTAATCAAAACCACCATATTTTTTTGGATCCC | Cloning (VQR vector) |
| QR-2-Rev | GAGTGGCATCTAAAACTTCTTTTGTAGACCTATATTGTTTACGATCAATTGTTGTATCAA | Cloning (E/VQR insert) |
| E-2-Fwd | GACTGGGATCCAAAAAAATATGGTGGTTTTGAAAGTCCAACGGTAGCTTATTCAG | Cloning (EQR insert) |
| V-2-Fwd | GACTGGGATCCAAAAAAATATGGTGGTTTTGTTAGTCCAACGGTAGCTTATTCAG | Cloning (VQR insert) |
